# Coordinated Regulation of Cdc42ep1, Actin, and Septin Filaments during Neural Crest Cell Migration

**DOI:** 10.1101/2022.12.04.519046

**Authors:** Mary Kho, Siarhei Hladyshau, Denis Tsygankov, Shuyi Nie

## Abstract

The septin cytoskeleton has been demonstrated to interact with other cytoskeletal components to regulate various cellular processes, including cell migration. However, the mechanisms of how septin regulates cell migration are not fully understood. In this study, we use the highly migratory neural crest cells of frog embryos to examine the role of septin filaments in cell migration. We found that septin filaments are required for proper migration of neural crest cells by controlling both the speed and the direction of cell migration. We further determined that septin filaments regulate these features of cell migration by interacting with actin stress fibers. In neural crest cells, septin filaments co-align with actin stress fibers, and the loss of septin filaments leads to impaired stability and contractility of actin stress fibers. In addition, we showed that a partial loss of septin filaments leads to drastic changes in the orientations of newly formed actin stress fibers, suggesting that septin filaments help maintain the persistent orientation of actin stress fibers during directed cell migration. Lastly, our study revealed that these activities of septin filaments depend on Cdc42ep1, which co-localizes with septin filaments in the center of neural crest cells. Cdc42ep1 interacts with septin filaments in a reciprocal manner, with septin filaments recruiting Cdc42ep1 to the cell center and Cdc42ep1 supporting the formation of septin filaments.

## INTRODUCTION

Septin filaments are the fourth component of the cytoskeleton after microfilaments (a.k.a. actin filaments), microtubules, and intermediate filaments. Septin filaments are assembled by a highly conserved family of GTP-binding proteins that were first discovered for their involvement in budding yeast morphogenesis (Hartwell et al., 1970). In humans, there are 13 known septins (Sept1-12 and Sept14), which are divided into four subgroups based on sequence similarity: Sept2, Sept3, Sept6, and Sept7 (Kinoshita, 2003; Martinez and Ware, 2004). The septin subunits assemble into non-polar hetero-hexamers and hetero-octamers through interactions between G (the GTP-binding) - G interfaces and NC (the N-/C-terminal) - NC interfaces (Sirajuddin et al., 2007). Septin heteromers further organize to form filaments, rings, and other higher-order structures, which have been observed to associate with the plasma membrane, actin filaments, and microtubules across different cellular processes (Marquardt et al., 2019; Mostowy and Cossart, 2012; Pous et al., 2016). During initial bud formation in *Saccharomyces cerevisiae*, septins form an hourglass-like structure that scaffolds actin polymerization and remodeling proteins in the nascent bud (Marquardt et al., 2019; Spiliotis and Nakos, 2021). Septins also recruit Myosin II for the formation of the actomyosin contractile ring during cytokinesis initiation (Feng et al., 2015; Garabedian et al., 2020). Recently, additional activities of septin filaments have been identified. In epithelial cells, septins are found to associate with microtubules to promote the formation of the apical microtubule network and the establishment of apicobasal polarity (Bowen et al., 2011). The septin-microtubule association has also been shown to play a role in neuronal dendrite and axon branching and in spermatozoa morphogenesis (Ageta-Ishihara et al., 2013; Kuo et al., 2013). In recent years, septin filaments have also been implicated in metastatic cancer cell migration and invasion (Fan et al., 2021; Farrugia et al., 2020; Xu et al., 2018; Zeng et al., 2019; Zhang et al., 2020). However, relatively little is known about the mechanism of how septin filaments regulate cell migration and how septin filaments are regulated during cell migration.

Neural crest cells represent a valuable model for studying directed cell migration. Born at the neural plate border, neural crest cells migrate long distances throughout the vertebrate embryo to give rise to a large variety of tissues and cell types, including craniofacial bones and cartilages, smooth muscles, and glial cells (Bronner and LeDouarin, 2012; Dupin et al., 2006). Interestingly, many signaling pathways and transcription factors involved in neural crest cell migration are also shared by metastatic cancer cells (Gallik et al., 2017). Therefore, regulators of cancer cell migration may also regulate neural crest cell migration, and knowledge gained from studying neural crest cell migration can contribute to a better understanding of the progression to cancer metastasis. Currently, the best-understood cytoskeletal components involved in neural crest cell migration are actin filaments. The Rho family of small GTPases plays important roles in regulating actin filaments during neural crest cell migration. Rac1 and Cdc42 are activated at the cell front and promote actin polymerization, resulting in protrusive activity (Chen et al., 2017; Ridley, 2011). At the trailing edge, RhoA is activated to regulate the contraction of myosin-associated actin stress fibers so that the cell rear is pulled towards the cell front (Matthews et al., 2008). But whether septin filaments also play a role in neural crest cell migration is still unknown.

In this study, we investigated the role of septin filaments in neural crest cell migration. We found that septin filaments are required for neural crest cell migration both *in vivo* and *in vitro*. We demonstrated that septin affects neural crest cell migration by regulating actin stress fibers. When the assembly of septin filaments was inhibited, the length and contractility of actin stress fibers were significantly reduced. Furthermore, our results revealed that septin interacts with Cdc42ep1 in neural crest cell migration. We have previously shown that Cdc42ep1 localized to both the cell periphery and center in neural crest cells. At the cell periphery, Cdc42ep1 interacts with Cdc42 to regulate actin dynamics and cell protrusions (Cohen et al., 2018). Here, we show that Cdc42ep1 interacts with septin filaments at the cell center. Our results indicate that localization of Cdc42ep1 at the cell center depends on septin filaments, while the organization of septin filaments is supported by Cdc42ep1. Our work also demonstrates that septin-Cdc42ep1 interaction regulates the directional migration of neural crest cells by regulating cell contraction along the direction of cell migration.

## RESULTS

### Septin filaments are required for neural crest cell migration

To determine whether septin filaments are required for neural crest cell migration, we disrupted septin filament formation in frog neural crest cells by knocking down the most indispensable subunit of septin filaments, Septin7. We used two methods to knock down Septin7, a translation-blocking morpholino (MO) and CRISPR-mediated genome editing. Migration of cranial neural crest cells after Septin7 knockdown was assessed by *in situ* hybridization against neural crest marker gene *Sox10*. Our results showed that both Septin7 knockdown by morpholino and Septin7 mutation by genome editing (Supplement al Table 1) led to apparent defects in neural crest cell migration (Figure 1A). While in control embryos or in the contralateral side of Septin7-MO injected embryos, neural crest cells migrated extensively into the developing branchial arches, their migration distance in CRISPR-targeted embryos or in the Septin7-MO injected side was significantly reduced. In parallel, we used a chemical inhibitor forchlorfenuron (FCF) (Hu et al., 2008) that inhibits septin filament formation in neural crest cells (Supplemental Figure S1A). Strong inhibition of neural crest cell migration was observed when 200μM of FCF was added to the embryo medium. As a negative control, we used ethanol, the solvent in which FCF was dissolved. The dose of ethanol used in our studies does not lead to any migration defect of neural crest cells (Supplemental Figure S1). The relative distance of neural crest migration across the entire dorsal-ventral axis of the embryos was quantified and summarized in Figure 1B. The three methods of septin inhibition decreased the relative distance of neural crest cell migration from 85% to 67%, 58%, and 57%, respectively. We also counted the number of distinct migratory streams of cranial neural crest cells. While control neural crest cells segregated into 2.8 migratory streams on average, Septin7-MO, Septin7 CRISPR, and FCF-receiving neural crest cells segregated into 1.7, 2.2, and 1.6 streams on average, respectively. The change in both the migration distance and the number of migratory streams was statistically significant (student *t*-test, p-value<0.001). When Septin7 RNA without the 5’UTR, which is the recognition site for Septin7-MO, was co-injected with Septin7-MO, the migration defects were efficiently rescued. Both the migration distance and the number of migratory streams were restored to comparable levels as in control embryos (81% of the D-V axis and 2.8 streams). These results demonstrate that septin filaments are required for the proper migration of neural crest cells in frog embryos.

**Figure 1.**
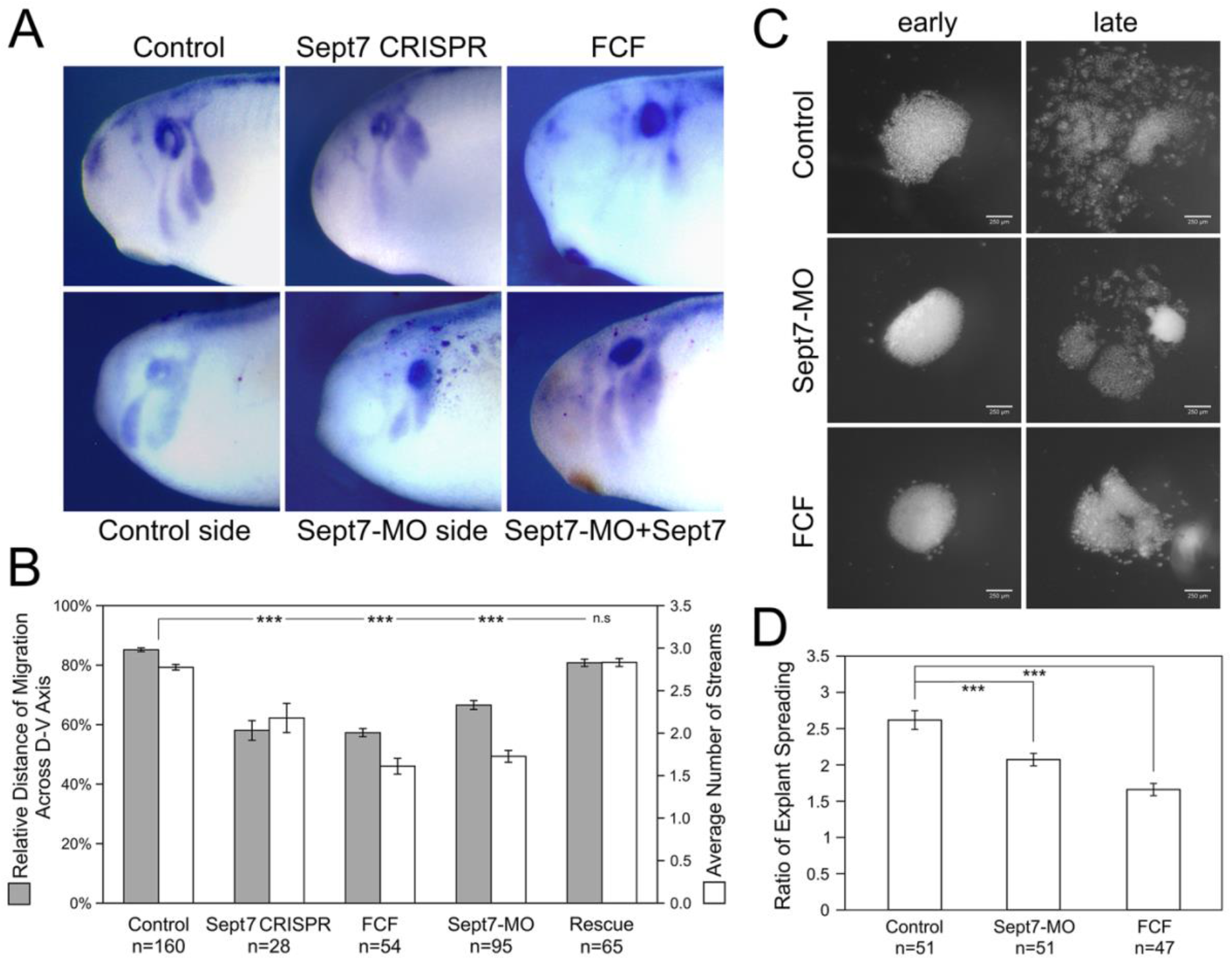
Loss of septin filaments leads to defective neural crest cell migration. A) Sox10 in situ hybridization on control and experimental embryos. Septin inhibition by CRISPR-mediated genome editing on Septin7, Septin7 translation blocking morpholino (10ng), and septin inhibitor forchlorfenuron (FCF, 200μM) all lead to obvious defects in neural crest cell migration. Septin7 RNA can rescue neural crest migration defects caused by Septin7-MO. Septin7-MO and RNA were co-injected into one side of the cleavage stage embryos together with the lineage tracer nuclear beta-Gal, which was detected by enzymatic reactions (shown as dark red dots). B) Relative distance of cranial neural crest cell migration across the dorsal-ventral axis of the embryo and the number of migratory streams were measured/counted and plotted. Septin7 CRISPR, Septin7-MO, and FCF treatment all led to a significant reduction in the migration distance and the number of migratory streams. Septin7 rescued the migration defects effectively. C) Spreading of neural crest explants on fibronectin. Images were taken 2 hours and 16 hours after dissection (around stage 17 and 24). D) The degree of explant spreading was calculated by comparing the area of explants at the early and late stages. Both Septin7-MO and FCF decreased the ratio of explant spreading significantly. ***, p<0.001.

To gain a better understanding of how septin filaments regulate neural crest cell migration, we isolated cranial neural crest tissue from late gastrula stage embryos, cultured them on fibronectin-coated cover glass, and observed their migration directly. We compared the migratory behavior of neural crest explants from control embryos, Septin7-MO injected embryos, and control explants cultured in a medium containing 200μM of FCF. First, the migration of neural crest explants was assessed by the ratio of explant spreading over time (from roughly stage 17 to stage 24). As shown in Figure 1C-D, septin inhibition slowed down neural crest explant spreading. While control explants had spread extensively with neural crest cells breaking away from the coherent cell sheet and migrating as individual cells or small cell clusters, most of Septin7-MO or FCF receiving cells were still in the collective migration phase and had migrated shorter distances. We measured the areas of neural crest explants at the early and late stages and compared the ratios of these areas. Septin7-MO and FCF reduced the ratio of explant spreading from 2.6 to 2.1 and 1.7, respectively. This reduction was statistically significant with p-values<0.001.

The defects in neural crest migration and explant spreading could result from decreased migration speed or failure to maintain persistent migration. To further elucidate how septin filaments regulate neural crest cell migration, we next tracked migrating neural crest cells in explant culture using the nuclear marker H2b-EGFP. The trajectories of cells were followed for 4 hours (see Supplemental Movies 3 and 4). Cell migration speed and persistence (directionality ratio) were calculated and plotted in Figure 5. Our results show that 10ng of Septin7-MO significantly reduced both the cell migration speed and the persistence of neural crest cell migration. These results indicate that septin filaments play important roles in neural crest cell migration and the loss of septin filaments leads to impaired directional migration of neural crest cells.

### Septin filaments co-align with actin stress fibers in neural crest cells and regulate the stability and contractility of actin stress fibers

To further determine how septin regulates neural crest cell migration, we examined its interaction with actin filaments. Actin is the key part of force-generating machinery that powers the locomotion of most cells. Septin filaments have been reported to associate with actin filaments in different cancer cells (Farrugia et al., 2020; Zeng et al., 2019). To determine whether septin filaments interact with actin filaments in neural crest cell migration, we first examined the subcellular localization of septin filaments and actin filaments in migrating neural crest cells. Immunofluorescence experiments were performed on neural crest explants against Septin7 (Invitrogen) and actin filaments (phalloidin, Invitrogen). Subcellular localization of the two types of filaments was visualized by confocal microscopy. As shown in Figure 2A and Supplemental Figure S3, Septin7 was observed near the plasma membrane at locations where there is inward membrane curvature and in bundles of filaments near the ventral surface where they co-align with actin stress fibers.

**Figure 2.**
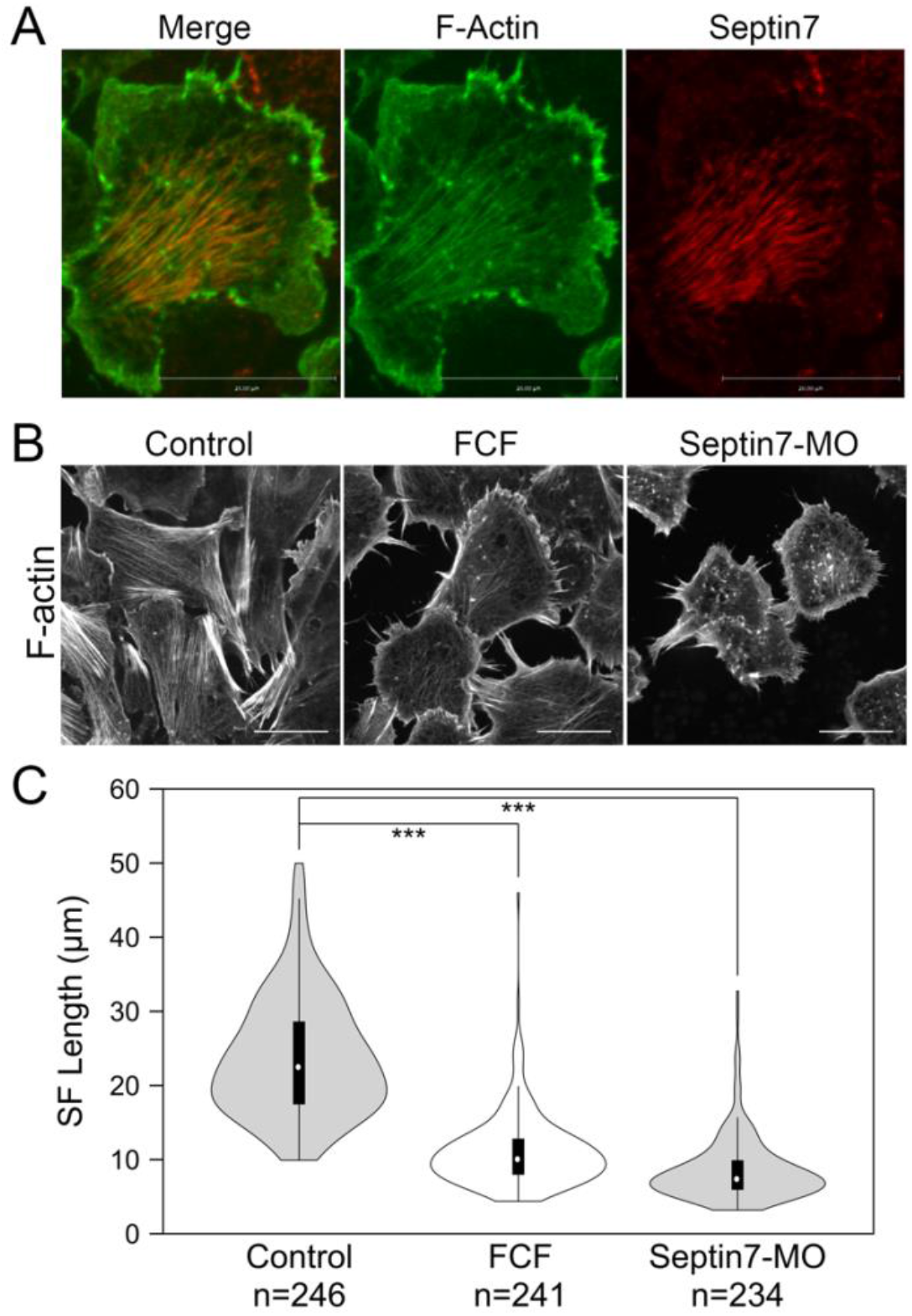
Septin filaments co-align with ventral actin stress fibers and is required for proper organization of actin stress fibers. A) Septin filaments co-align with actin stress fibers in neural crest cells. Septin7 antibody and phalloidin were used to visualize septin filaments and actin filaments, respectively. B) Inhibition of septin filaments by both Septin7-MO and FCF treatment leads to defects in actin stress fiber formation. Actin filaments were visualized by live imaging against GFP-Utrophin. C) The length of actin stress fibers was measured and plotted in violin plots. Both FCF and Septin-MO reduced the length of actin stress fibers significantly. ***, p<0.001. Scale bar = 20μm.

Co-alignment between septin filaments and actin stress fibers suggests that septin filaments may regulate the formation or contractility of actin stress fibers. We next examined whether septin filaments play a role in the formation of actin stress fibers. Neural crest explants were prepared from embryos injected with GFP-Utrophin RNA so that actin stress fibers could be visualized. The assembly of septin filaments was disrupted by co-injecting Septin7-MO or by adding FCF in the culture medium. The effects of such septin disruption on actin stress fibers were analyzed by live imaging. When the formation of septin filaments was inhibited, the formation of actin stress fibers was impaired as well. Long actin stress fibers were rarely observed. Instead, actin formed shorter fibers or aggregates that were scattered in the cell (Figure 2B). We measured the length of actin stress fibers using the line selection tool in ImageJ. The results plotted in Figure 2C show a significant decrease in the length of actin stress fibers from an average of 24 μm in control cells to below 12 μm in FCF-treated or Septin7-MO injected cells.

The decrease in the length of actin stress fibers could result from defects in their assembly or stability. To determine whether septin filaments play a role in the assembly of actin stress fibers, we performed an actin disassembly-reassembly assay. Actin stress fibers were first removed by treating neural crest cells with actin polymerization inhibitor Latrunculin A (Lat A, Tocris Bioscience). Lat A has been shown to not only inhibit the formation of new actin filaments but also promote the depolymerization of existing actin filaments (Fujiwara et al., 2018). We found that 1 hour of Lat A treatment (100nM) is sufficient to remove all actin stress fibers as well as apparent actin filaments inside the neural crest cells. Disassembled actin formed aggregates that were scattered throughout the cell. Then, we washed the cells thoroughly to remove Lat A and allowed actin stress fibers to form again. The recovery of actin stress fibers was compared between cells with and without FCF added into the medium after Lat A washout (Supplemental Movies 1-2). As shown in Figure 3A, fine actin filaments started to appear in control cells after 2-4 minutes, and they assembled into thick actin stress fibers by 6-12 minutes. We did not notice a significant difference in the recovery of actin stress fibers when FCF was present. It took 4.375 minutes on average for cells to assemble actin filaments and 8.375 minutes on average for actin stress fibers to become prominent in FCF-treated cells. However, without the support of septin filaments, the actin stress fibers were soon broken apart and reduced to small aggregates. These results indicated that in neural crest cells, septin filaments are not required for the assembly of actin stress fibers but instead play an important role in the stability of actin structures.

**Figure 3.**
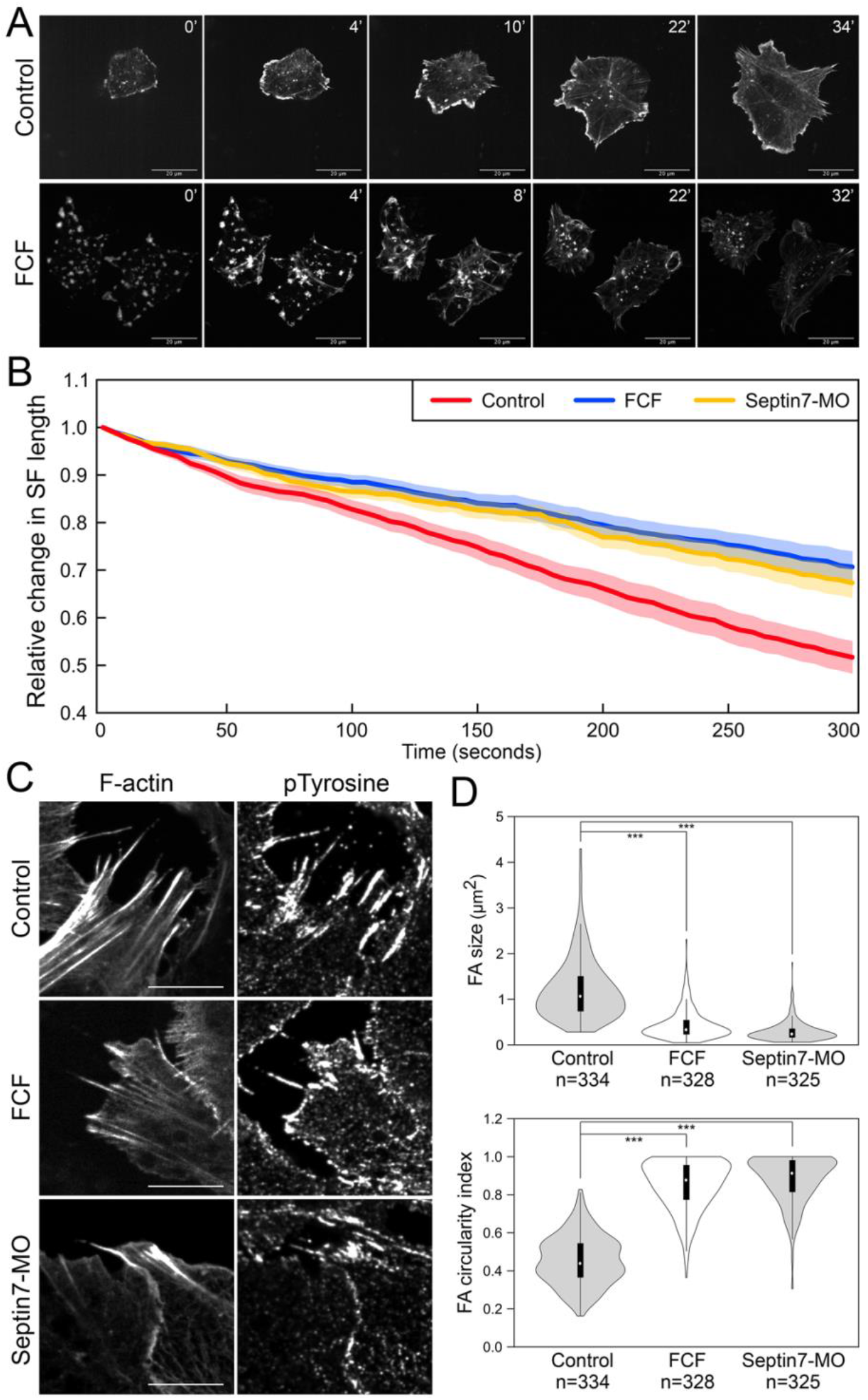
Septin filaments regulate the stability and contractility of actin stress fibers. A) Recovery of actin stress fiber in control and FCF treated neural crest cells. Actin stress fibers were removed in neural crest cells by Latrunculin A treatment and their reassembly were documented by time-lapse imaging. While actin stress fibers were reassembled in FCF-treated cells, they tended to disappear over time. Scale bar = 20μm. B) The contractility of actin stress fibers was quantified by measuring the dynamic change of their lengths. The relative changes in the length of 20 actin stress fibers per condition were plotted. Both Septin7-MO and FCF treatment significantly decreased the speed of actin stress fiber shortening (p<0.001). C) Septin inhibition disrupted the formation of focal adhesions. Separate images of focal adhesion (immunofluorescence against pTyr) and actin filaments (GFP-Utrophin) were shown. Scale bar = 10μm. D) Analysis of focal adhesion size and circularity. Both methods of septin inhibition reduced focal adhesion size and elongation significantly. *** p<0.001.

When septin filaments were disrupted, a small number of actin stress fibers were still present. We further examined whether the contractility of these stress fibers was affected by the loss of colocalization with septin filaments. We observed the dynamics of actin stress fibers and measured their length over time. Since many actin stress fibers disappeared quickly after their assembly, only stress fibers that lasted 5 minutes were used for our measurement. The changes in the lengths of 20 actin stress fibers were plotted in Figure 3B for each condition. Compared to stress fibers in control neural crest cells, which shorten rapidly during contraction, actin stress fibers in Septin7-MO injected or FCF treated cells contract significantly slower. These observations suggest that the contractility of actin stress fibers was impaired by the loss of septin filaments.

Since focal adhesions are linked to actin stress fibers and change their shapes in response to the mechanical tension that they receive through actin stress fibers, we next assessed the contractility of actin stress fibers by measuring the shape of focal adhesions. Focal adhesions were visualized by either immunofluorescence against pTyr (Figure 3C) or by live imaging of the fusion protein EGFP-FAK (Supplemental Figure S2). While elongated focal adhesions were observed at the end of actin stress fibers in control neural crest cells, focal adhesions became much shorter or reduced to small round dots when septin assembly was inhibited. The size and shape of focal adhesions were quantified by the oval area selection tool in ImageJ. Loss of septin filaments resulted in a decrease in the size of focal adhesions from 1.2 μm^2^ to 0.4 and 0.3 μm^2^, while the circularity index of focal adhesions increased from 0.45 to 0.85 and 0.88, respectively. These results imply a significant decrease in the tension on focal adhesions due to a decrease in the contractility of stress fibers. In summary, septin filaments play important roles in both the stability and the contractility of actin stress fibers. Therefore, septin filaments likely regulate neural crest cell migration by regulating actin stress fibers.

### Cdc42ep1 colocalizes with septin filaments and regulates the assembly of septin filaments

It has been reported in cancer cells and in endothelial cells that Cdc42 effector proteins interact with septin filaments to regulate cell migration (Calvo et al., 2015; Liu et al., 2014). We have previously reported that in cranial neural crest cells, Cdc42ep1 is highly expressed in neural crest cells and interacts with Cdc42 at the cell protrusions to regulate actin dynamics and cell motility (Cohen et al., 2018). Here, we ask whether Cdc42ep1 also interacts with septin filaments in neural crest cells. First, we analyzed colocalization between Cdc42ep1 and Septin7 in neural crest explants. Cdc42ep1 was visualized by live imaging using fluorescent fusion protein Cdc42ep1-EGFP, and Septin7 was detected by immunofluorescence (Figure 4A). Our results showed that Cdc42ep1 colocalizes with Septin7 with a colocalization index of 0.74. In addition to colocalizing with septin filaments of different lengths, Cdc42ep1 also colocalizes with septin rings, which were predicted to form when septins dissociate from actin filaments (Kinoshita et al., 2002). On the contrary, a mutant form of Cdc42ep1, Cdc42ep1 (GPS) with reported defective binding to septin (Joberty et al., 2001), distributed in a diffused manner in neural crest cells and failed to colocalize with septin filaments. The correlation coefficient between Cdc42ep1 (GPS) and Septin7 has a significantly lower value of 0.21. These results suggest that Cdc42ep1 is recruited to its cell center localizations by septin filaments.

**Figure 4.**
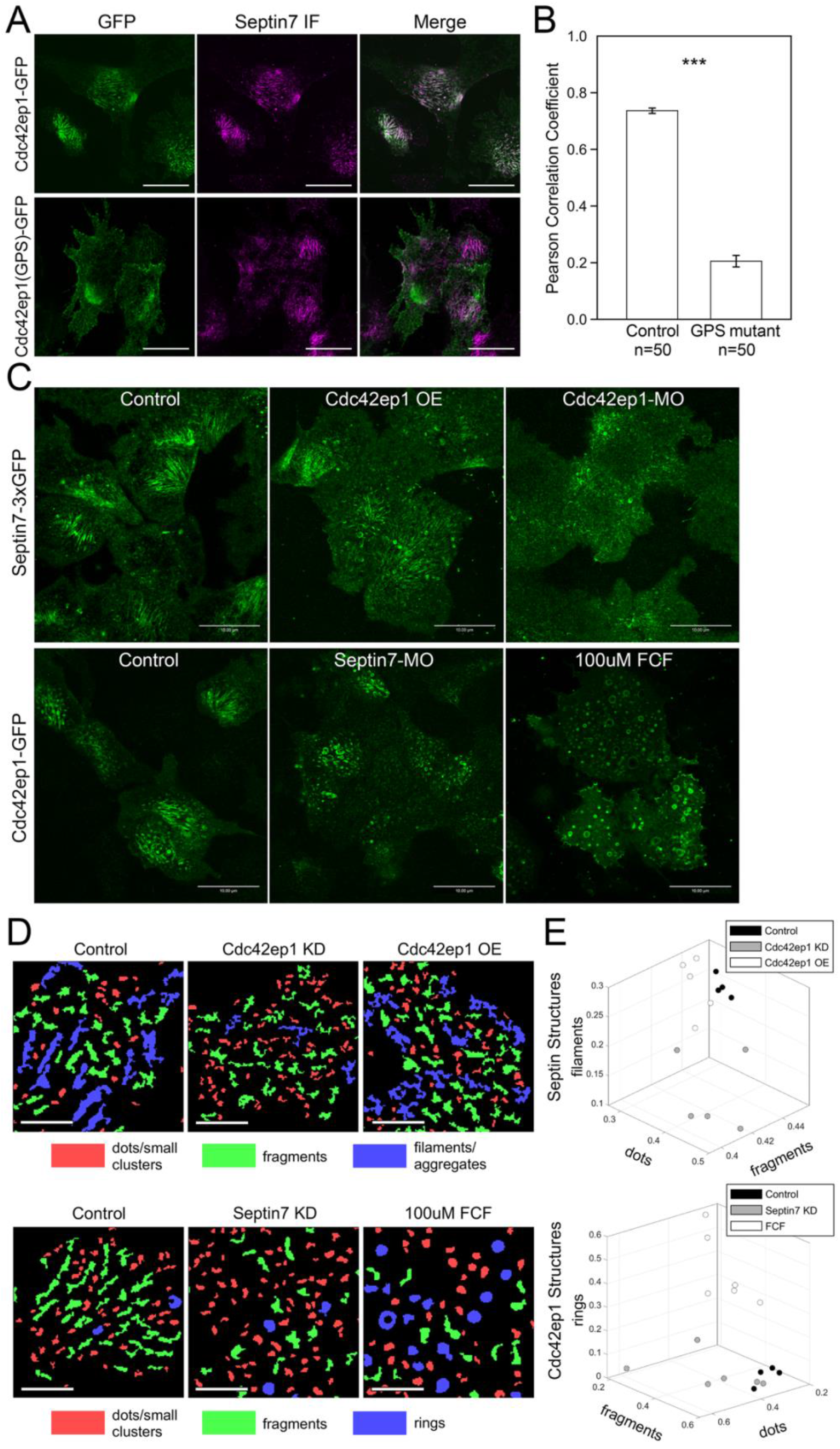
Cdc42ep1 colocalizes with septin filaments and interacts with septin filaments in a reciprocal manner. A) Wild type Cdc42ep1, not the septin binding defective mutant form of Cdc42ep1, colocalizes with septin filaments in neural crest cells. B) Correlation coefficient of Cdc42ep1 and septin7. There is a significant decrease in the correlation coefficient between septin binding defective Cdc42ep1 and septin filaments. C) Misexpression of Cdc42ep1 disrupted the organization of septin filaments, while septin inhibition affected the cell center localization and organization of Cdc42ep1. Scale bar = 10μm. D) Septin and Cdc42ep1 structures were classified into three major groups (see Methods and Supplemental Figure S4-5). The three groups of structures were pseudo-color-coded in example images. Scale bar = 5μm. E) The area fraction of structures in the three identified classes was plotted. Phenotypic distinction between different experimental manipulations were observed, demonstrating distinct effect from different experimental treatments.

**Figure 5.**
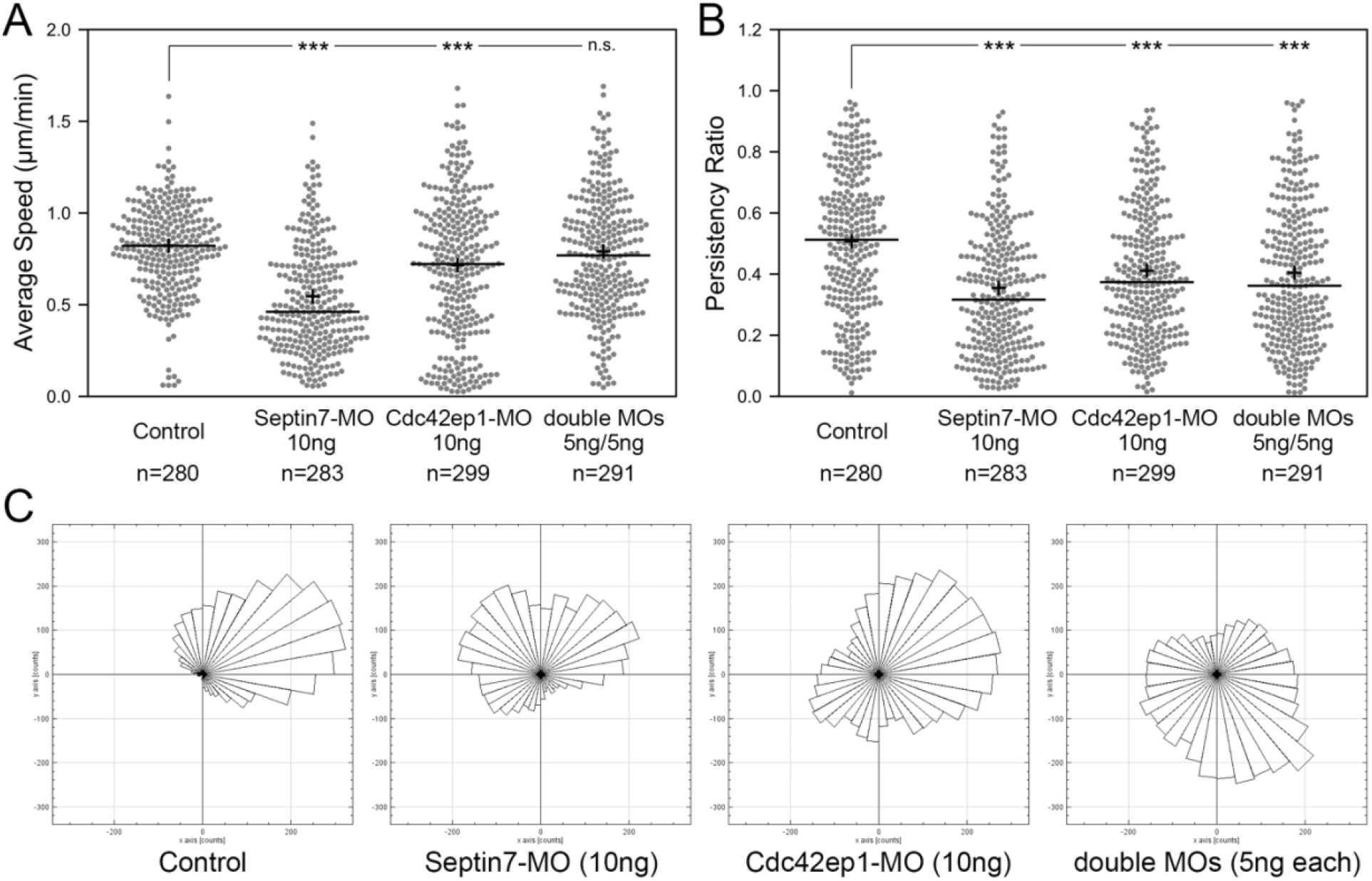
Septin filaments and Cdc42ep1 cooperate in regulating the directional migration of neural crest cells. A) The effect of Septin7 knockdown, Cdc42ep1 knockdown, and double knockdown on the speed of neural crest cell migration. B) The effect of Septin7 knockdown, Cdc42ep1 knockdown, and double knockdown on the persistence of neural crest cell migration. +, mean. —, median. ***, p<0.001. C) Rose plots of representative neural crest explants demonstrating the directional migration of neural crest cells.

Colocalization between Cdc42ep1 and Septin7 suggests that there is a functional interaction between the two. Next, we examined how Cdc42ep1 regulates the assembly of septin filaments. Cdc42ep1 was either overexpressed or knocked down (by a translation-blocking morpholino), and the effects of these perturbations on the formation of septin filaments (as reflected by Sept7-3xGFP) are shown in Figure 4C. When Cdc42ep1 was overexpressed, the formation of long septin filaments was moderately reduced with a concurrent increase of septin aggregates or rings. When Cdc42ep1 was knocked down, the assembly of septin filaments at the cell center was dramatically disrupted, resulting in a large number of short septin fragments. The organization of different septin structures was quantified with an automated classification pipeline (see Methods), with the results summarized in Figure 4D-E. Our results show that the expression level of Cdc42ep1 impacted the organization of septin structures. Overexpression of Cdc42ep1 tends to increase the area fraction of septin aggregates. Knockdown of Cdc42ep1 resulted in significantly more diffused septin puncta at the expense of septin filaments and large aggregates. These results indicate that an optimal level of Cdc42ep1 is critical for the proper organization of septin filaments.

When changes in Cdc42ep1 expression disrupted the organization of septin filaments, the organization of actin stress fibers was also impaired (Supplemental Figure S3). When Cdc42ep1 was overexpressed, we observed an increase in septin aggregates. Concurrently, actin stress fibers were often thinner or disorganized. When Cdc42ep1 was knocked down, septin filaments appeared fragmented, and the localization of Septin7 became diffused. At the same time, we observed much fewer actin stress fibers. However, despite the defects in septin filament assembly and actin stress fiber formation, Septin7 molecules still resided closely to actin structures, suggesting that the association between septin and actin does not depend on Cdc42ep1.

Since Cdc42ep1 colocalizes with and is recruited by septin filaments, we next determined how defective septin filament formation affects the organization of Cdc42ep1. Assembly of septin filaments was inhibited by either Septin7-MO or FCF, and the subcellular localization of Cdc42ep1 was examined by live imaging using Cdc42ep1-GFP expressed in the cell (Figure 4C). In control neural crest cells, Cdc42ep1 assembles into filamentous structures, arcs, and occasionally rings, similar to the subcellular organization of septins (Figure 4A). When septin filament formation was disrupted, the filamentous appearance of Cdc42ep1 was gone. Cdc42ep1 aggregated along fragmented structures and rings that were distributed throughout the cell. The organization of Cdc42ep1 structures was also quantified, and the result was summarized in Figure 4D-E. Both methods of septin inhibition disrupted the filamentous arrangement of Cdc42ep1, with Septin7 knockdown leading to the formation of fragmented Cdc42ep1 aggregates and an increased number of puncta. FCF treatment led to the formation of circular structures of intermediate and large sizes. This result indicates that in addition to the role of Cdc42ep1 in supporting the formation of septin filaments, septin filaments also play a reciprocal role in the subcellular organization of Cdc42ep1.

### Septin and Cdc42ep1 cooperatively regulate directional migration of neural crest cells

Since both Cdc42ep1 and septin filaments regulate neural crest cell migration and there are reciprocal interactions between the two, we sought to examine how they cooperate during neural crest cell migration. 10ng of Cdc42ep1-MO, 10ng of Septin7-MO, or 5ng of both Cdc42ep1-MO and Septin7-MO were injected into frog embryos, together with the nuclear tracer H2b-EGFP. The migration of neural crest cells in explant culture is shown in Supplemental Movies 3-6, and the analysis of neural crest cell trajectories is shown in Figure 5. While 10ng of Cdc42ep1-MO or Septin7-MO led to a significant decrease in both the speed and persistence of neural crest cell migration, the combination of half dosage of both morpholinos only affected the persistence of neural crest cell migration, but not the speed of cell migration (Figure 5A-B). Rose plots of cell migration further demonstrate that the combined loss of Cdc42ep1 and Septin7, despite at lower dosage, dramatically abolished the directional migration of neural crest cells (Figure 5C). Although having been isolated from the embryo, neural crest cells still maintained some of their original polarity and migrated in one direction as a group. This unidirectional collective motion was lost when Cdc42ep1 or Septin7 was knocked down, demonstrating a critical role for Cdc42ep1 and septin filaments in maintaining the directional migration of neural crest cells.

To better understand how Cdc42ep1 and septin filaments maintain directional migration of neural crest cells, we re-examined the dynamics of actin filaments during neural crest cell migration (Supplemental Movies 7-10). In control cells, actin filaments were present in both membrane protrusions and stress fibers. The actin stress fibers were aligned parallel to the direction of cell migration and contracted efficiently to pull the cell forward. When actin stress fibers were disassembled, new actin stress fibers were assembled with a similar orientation, likely using the more stable septin filaments as templates. When Cdc42ep1 or Septin7 was knocked down by 10ng of morpholino, the assembly of actin protrusions and stress fibers was suppressed. When Cdc42ep1 was knocked down, neural crest cells no longer made lamellipodia but moved with membrane blebs, as described previously (Cohen et al., 2018). When Septin7 was knocked down, lamellipodia and membrane blebs coexisted, but neither was able to lead cell migration. Under both knockdown conditions, actin stress fibers were not evident, and cells kept changing their direction of migration. When both Cdc42ep1 and Septin7 were partially knocked down by 5ng of morpholino, both actin protrusions and actin stress fibers were observed. However, after the disassembly of existing actin stress fibers, new actin stress fibers were assembled in a different orientation, leading to a change in the direction of migration. This observation provides a plausible explanation for the defective directional migration of neural crest cells: the loss of Cdc42ep1 and septin filaments leads to the destabilization of actin stress fibers and the loss of continuity of stress fiber orientation during the disassembly-assembly cycle. This, in turn, abolished the persistent alignment of actin stress fibers and disrupted the directional contraction of cells along their path of migration. This study, therefore, demonstrates that the cooperation between Cdc42ep1 and septin filaments regulates the directional migration of neural crest cells by enabling the persistence of actin stress fiber orientation.

## DISCUSSION

In this study, we investigated the role of septin filaments in neural crest cell migration. We showed that septin filaments are required for neural crest cell migration and control both the speed of cell migration and the persistence of cell migration. We further demonstrated that septin filaments regulate the speed of neural crest cell migration by regulating the contractility of actin stress fibers. Septin also regulates the persistence of neural crest cell migration by maintaining the persistent orientation of actin stress fibers. These activities of septin filaments depend on Cdc42ep1. Cdc42ep1 is recruited to the cell center by septin filaments and supports the formation of septin filaments. Cdc42ep1 plays this role in addition to its previously reported role, in which it interacts with Cdc42 at the cell protrusions to promote their extension (Cohen et al., 2018). Together, our works suggest that Cdc42ep1 plays a critical role by coordinating the protrusive front and retractive rear of neural crest cells, thus regulating their directed migration.

### Interaction between septin filaments and actin filaments

Our results suggest that in neural crest cells, septin filaments do not regulate actin polymerization or actin stress fiber assembly but instead regulate the stability of actin stress fibers. This result is different from the reported function of Septin6 in chick dorsal root ganglia neurons, where Septin6 colocalizes with cortactin and promotes actin polymerization through Arp2/3 (Hu et al., 2012). The different activity of septins likely results from different interacting partners of septins in different cells. We showed that septin filaments promote the stability of actin stress fibers in neural crest cells, and this function is likely mediated by additional actin-binding proteins. Septin9 is currently the only septin subunit that has been shown to bind directly to actin filaments (Smith et al., 2015). Currently, we do not know whether Septin9 mediates the interaction between septin filaments and actin filaments in neural crest cells. Another potential mediator is Myosin II. It has been reported that septins directly interact with Myosin II during cytokinesis and in cell migration (Feng et al., 2015; Garabedian et al., 2020; Joo et al., 2007; Zeng et al., 2019). The septin-Myosin II interaction not only regulates the activity of Myosin II, therefore regulating the contractility of actomyosin, but also plays a part in the septin-actin association (Joo et al., 2007). It is possible that in neural crest cells, the interaction between septin filaments and Myosin II also mediates the association between septin filaments and actin filaments.

### Interaction between septin filaments and Cdc42ep1

We demonstrated that septin filaments and Cdc42ep1 regulate each other in a reciprocal manner. When Cdc42ep1 was overexpressed, we observed aggregates of septin molecules. When Cdc42ep1 was knocked down, septin filaments disassembled, and septins were distributed in a diffused manner. These results suggest that Cdc42ep1 plays an important role in the stabilization of septin filaments. Since Cdc42ep1 can directly bind to septins, it may stabilize septin filaments as a side binder. In MDCK cells, another Cdc42 effector protein, Cdc42ep5, has been shown to regulate septin filament assembly and organization (Joberty et al., 2001). Similarly, Cdc42ep3 has been shown to regulate the organization of septin filaments in cancer-associated fibroblast cells (Calvo et al., 2015). In breast cancer cells, Cdc42ep3 or Cdc42ep5 have also been implicated in promoting the septin-actin association (Salameh et al., 2021; Zeng et al., 2019). When these Cdc42eps were reduced, septins dissociated from actin filaments and localized with microtubules (Salameh et al., 2021). In neural crest cells, when Cdc42ep1 was reduced, the assembly of both septin filaments and actin filaments was reduced, but the remaining septin filaments were still localized next to actin filaments (Supplemental Figure S3). Whether the dissociated septins are associated with microtubules remains to be elucidated.

When septin filament assembly was inhibited by Septin7-MO or FCF, the organization of Cdc42ep1 was dramatically affected. Cdc42ep1 was no longer organized along filamentous structures and instead formed aggregates or rings of different sizes. When Septin7 was knocked down, the Cdc42ep1 rings or aggregates were generally small in size. However, upon FCF-mediated septin inhibition, Cdc42ep1 made ring structures with a wide range of sizes. The diameter of these ring structures ranges from 0.49μm to 1.35μm, exceeding the range of typical septin rings (0.66-0.68μm) (Kinoshita et al., 2002). There are two possible explanations for such a difference. First, Cdc42ep1 may have a higher affinity to Septin7 compared to other septin molecules. This is the case for Cdc42ep5, which binds specifically to Septin6-Septin7 heterodimers (Sheffield et al., 2003). When Septin7 expression was reduced, Cdc42ep1 became more diffused. When Septin7 was present, but the organization of septin filaments was defective (in FCF treatment), Cdc42ep1 surrounded these septin7-containing structures and formed rings of different sizes. Alternatively, septins may form different structures under the two conditions. In the absence of Septin7, the remaining septin subunits may form oligomers of similar sizes. Under FCF treatment, however, septins may form aggregates of different sizes, which recruited Cdc42ep1 to form different-sized rings.

### Septin-Cdc42ep1 in neural crest migration

We found that when the expression level of Septin7 and Cdc42ep1 was moderately reduced, the speed of neural crest cell migration was not affected, but the directional persistence of cell migration was lost. This is different from the effect of a more complete knockdown of either Septin7 or Cdc42ep1, which leads to significant defects in both cell migration speed and persistence. We have concluded that this phenomenon results from different degrees of defects in actin stress fibers, which are critical for both cell motility and the direction of cell migration. When there is a dramatic decrease in Septin7 or Cdc42ep1 expression, cells barely form any actin stress fibers. As a consequence, cells migrate much slower and lose their direction of migration. At partial knockdown of both proteins, the remaining Cdc42ep1 can still support the assembly of septin filaments by the smaller number of available Septin7, which stabilizes some actin stress fibers. However, when existing actin stress fibers were disassembled, new actin stress fibers were often formed with a new orientation, leading to a change in migration direction. Therefore, cells can still migrate but randomly change their direction of migration over time. The failure to maintain the orientation of actin stress fibers is likely due to the deficiency of septin filaments. While many septin filaments colocalize with actin stress fibers, there are also septin filaments that do not associate with actin filaments. They simply align with each other in the space between actin stress fibers. It is possible that they serve as scaffolds for the assembly of new actin stress fibers irrespective of other stress fibers. Since septin filaments are more stable compared to actin filaments (Hagiwara et al., 2011), after the disassembly of old actin stress fibers in control cells, the coaligned septin filaments are still present with a similar orientation and, therefore, can guide the assembly of new actin stress fibers. In summary, septin filaments play an important role in the alignment of actin stress fibers and the directional migration of neural crest cells. Whether septin filaments regulate other directional migration events through the same mechanism remains to be examined.

## METHODS

### Embryo manipulations

*Xenopus laevis* embryos were obtained and staged as described by Nieuwkoop and Faber’s Normal Table of *Xenopus laevis*. The embryos were microinjected with guide RNA and Cas12a (cpf1) nuclease at 1-cell stage, or with capped RNAs or morpholino oligomers (MO) at 8-cell stage into dorsal animal blastomeres. Septin7 guide RNA (5’-GGTAGTCTTCAAATTTACTGTC −3’) that recognizes the Septin7 locus on both Chromosome 6L and 6S near D116 and Cas12a nuclease (Alt-R Cas12a) were purchased from IDT DNA. 2μM of guide RNA and 2ng of nuclease were injected together into 1-cell stage embryos (total 10nl in volume). Septin7-MO (5’-TGCTGTAGAGTCAGTGCCTCGCCTT −3’) hybridizes to −26 to −12 position relative to the translational start site of *Xenopus* Septin7 (GenBank Accession No. NM_001092714), Cdc42ep1-MO (described in (Cohen et al., 2018)), and standard control MO (Gene Tools, Philomath, OR) were used in the study. The Septin7 fluorescent fusion construct was generated by inserting the Septin7 sequence into the pCS2+3xEGFP construct (gift from Dr. Ann Miller from University of Michigan) as previously described (Cohen et al., 2018). Septin7 without 5’UTR (therefore cannot be recognized by Septin7-MO) and Cdc42ep1(GPS) mutant that is defective for septin binding were generated by site-directed mutagenesis using full-length Septin7 and Cdc42ep1-EGFP as a template, respectively. RNAs for fluorescent fusion proteins, wild type and mutant Septin7, H2b-EGFP, EGFP-FAK, and nuclear beta-galactosidase (nβGal) were synthesized with linearized templates using SP6 polymerase (Ambion mMessage mMachine Kit). All experimental procedures were performed according to USDA Animal Welfare Act Regulations and have been approved by Institutional Animal Care and Use Committee, in compliance of Public Health Service Policy.

### Red-Gal staining, in situ hybridization, and immunofluorescence

For lineage tracing, embryos co-injected with nβGal were fixed at the desired stage for half an hour in the fixative MEMFA, and stained with the Red-Gal substrate (Research Organics) until they turned red. The embryos were refixed for 2 hours in MEMFA and stored in methanol before *in situ* hybridization was performed. Whole mount *in situ* hybridization was performed as previously described (Cohen et al., 2018). Antisense probes for Sox10 were synthesized with T7 RNA polymerase with linearized plasmids. Immunofluorescence analysis was performed on neural crest explants plated on fibronectin-coated cover slides. The explants were fixed in MEMFA for 20-30 minutes and rinsed with PBT (PBS with 0.1% Triton X-100). They were then blocked in 10% goat serum and incubated with primary antibodies (anti-phosphotyrosine antibody (4G10; EMD Millipore), 1:200; anti-Septin7 (Invitrogen), 1 μg/mL; anti-Cdc42ep1 (produced at BosterBio), 5 μg/mL). Alexa Fluor-488/555 conjugated secondary antibody was used at 1:400 and the explants were imaged using a Nikon Ti2-E microscope (60x 1.4 N.A. objective).

### Cranial neural crest explant culture and microscopy

Cranial neural crest (CNC) explants receiving different MOs or RNAs encoding fluorescent proteins were dissected from stages 13-15 embryos as previously described (Alfandari et al., 2003; Borchers et al., 2000; DeSimone et al., 2005). Explants were plated onto fibronectin (FN, 20 μg/ml in PBS)-coated dishes in Danilchik media. Cell migratory behaviors and protein localizations were recorded as indicated by time-lapse microscopy using a PerkinElmer or Nikon Spinning Disc confocal microscope. To inhibit septin filaments, FCF (Forchlorfenuron, Acros Organics, 100 μM) was added into the culture medium. For actin stress fiber disassembly-reassembly assay, explants were treated with Lat A (Latrunculin A, Tocris Bioscience, 100 nM) for an hour before Lat A was removed by extensive washing.

### Image processing and quantification

To quantify the length of actin stress fibers, the straight-line selection tool in Fiji (ImageJ) was used to trace and measure the length of individual actin stress fibers. The oval selection tool in Fiji was used to quantify the size and shape of focal adhesions. To quantify the colocalization between Cdc42ep1 and Septin7, images were first subjected to deconvolution by using the Diffraction PSD 3D and Iterative Deconvolve plugins in Fiji. Entire cells were traced as regions of interest for calculation of Pearson’s R value (no threshold) by the Coloc2 plugin in Fiji.

To track cell trajectories and quantify the speed, persistence, and direction of cell migration, the TrackMate plugin in Fiji was used to detect and track the migration trajectories of individual cells from movies of neural crest explants labeled with H2b-EGFP. Movies were captured at 5-minute intervals for a total of 4 hours. Only tracks that persisted for more than 5 frames (more than 25 minutes) were selected for analysis. To analyze the contraction of actin stress fibers, movies of GFP-Utrophin-labeled neural crest cells were taken at 5-second intervals for 5 minutes. Using the straight-line selection tool in Fiji, the change in the length of actin stress fibers over time was calculated.

### Classification pipeline for Septin and Cdc42ep1 spatial organization

All steps of the pipeline were performed using MATLAB scripts developed for this work. The input data for the analysis were 16-bit grayscale images of live cells. Because of the variability of the background intensity, threshold values for cell segmentation were chosen manually for each image (the threshold ranged between 12000 and 24000 for septin data and between 2000 and 5000 for Cdc42ep1 data). The binary cell masks were further processed by (1) filling holes, (2) applying morphological opening (erosion followed by dilation) with a disk-shaped structuring element of radius 7 pixels, and (3) filtering out all connected objects with an area less than 5000 pixels.

For segmenting septin and Cdc42ep1 structures within the cells, we first performed local background subtraction using a Gaussian smoothing kernel with a standard deviation of 5 pixels. This step allowed us to even the background across the image and include both bright and dim structures in our analysis (Supplemental Figure S4A). To obtain the binary masks of individual structures, we used the threshold of 2000 for septin data and 1500 for Cdc42ep1 data. Any objects outside of cell masks were excluded from further analysis. Objects with an area of fewer than 10 pixels or more than 400 pixels were also excluded.

For each binary object (representing septin or Cdc42ep1 clusters, aggregates, or other formations), we obtained boundary coordinates using the built-in MATLAB function “bwboundaries” (with ‘noholes’ option). The objects’ masks and their boundary coordinates were used to characterize each object with 72 measures (hereon referred to as features, Supplemental Figure S4B). The first eight features were computed with the built-in MATLAB functions “regionprops”, which include ‘Area’, ‘ConvexArea’, ‘Eccentricity’, ‘EquivDiameter’, ‘Extent’, ‘FilledArea’, ‘MajorAxisLength’, ‘MinorAxisLength’, ‘Perimeter’, and ‘Solidity’. The next four features represented the mean and standard deviation of the minimal distance from the object’s boundary points to the nearest neighboring object and the mean and standard deviation of the intensity values of pixels within each object. The next 30 features were computed as Fourier modes on the boundary coordinates as described in Alizadeh et al. (2019). Such truncated Fourier decomposition approximates the object’s shape with a smooth contour. On rare occasions, when the error in the boundary approximation (i.e., the mean distance between the original and approximated boundary points) was more than 0.1 pixel, the structure was excluded from the analysis. Using the absolute values of the complex-valued amplitudes provides a rotationally invariant shape characterization. Similarly, the minimal distance from the object’s boundary points to the nearest neighboring object (calculated with the built-in MATLAB function “bwdist”) was approximated with the Fourier decomposition, providing the remaining 30 features. All computed features were standardized by subtracting the mean and dividing by the standard deviation.

The dimensionality of the feature space was reduced by using 17 principal components that explain 90% of data variance. For classification, we used the DBSCAN algorithm with parameters ε=2 and the minimal number of points for the core identification equal to 3 (Supplemental Figure S4C). The algorithm identified nine classes (together containing 99.5% of the data points) for septin data and eight classes (together containing 98.5% of the data points) for Cdc42ep1 data. Our pipeline includes an optional step of class refinement, which was needed for Cdc42ep1 data to improve the separation of small elongated and round objects (including disconnected/fragmented rings, see Supplemental Figure S4D). Refinement was performed for 5 classes with the largest object areas. For each class, we identified the principal component that showed the most pronounced bimodality. For the two most populated classes (with the smallest object areas), we used the Gaussian Mixture Model (GMM) with the default parameters in the MATLAB implementation. For the remaining three classes with a significantly smaller number of data points, instead of GMM, we used simple thresholding of the selected PC value. Based on the obtained classification of all objects, we performed image-level quantification using the area fraction metric, which is the ratio of the total area of objects in one class to the total area of objects in all classes (Supplemental Figure S5A).

To provide an intuitive visual interpretation of our results, we reduced the number of classes to three (“dots/small clusters,” “fragments,” and “filaments/aggregates” for septin data, and “dots/small clusters,” “fragments,” and “rings” for Cdc42ep1 data). Class merging was performed based on the co-alignment of the unit vectors of the area fraction measures along the first two principal components (Supplemental Figure S5B, C).

### Genomic PCR

Genomic DNA was extracted from individual frog embryos at tailbud stages with 100ul of genomic DNA extraction buffer (50mM Tris pH 8.8, 1mM EDTA, 0.5% Tween-20, 200μg/ml protease K). 2μl of the genomic DNA was used as a template for PCR reaction. The PCR product was ligated into pCR™Blunt II-TOPO™ vector (ThermoFisher Scientific) and sequenced using SP6 as the primer. Primers for genomic PCR: F: 5’-TGGCACAAACTTCTTGCACTT −3’; R: 5’-TCCATTTATTTCACCTGTAAGCCAT −3’.

## Acknowledgements

This work was supported by the National Institutes of Health (R01GM136892) to S.N. and D.T., and the National Science Foundation (CMMI 1942561) to D.T.

## Author Contributions

M.K. designed and performed experiments, analyzed data, and contributed to the writing of the manuscript. S.H. performed the imaging processing and analysis in Figure 4. D.T. oversaw the imaging analysis done by S.H. and revised the manuscript. S.N. designed the studies, oversaw the experiments and analyses and prepared the manuscript.

