## Supplemental Material for "Coordinated Regulation of Cdc42ep1, Actin, and Septin Filaments during Neural Crest Cell Migration"

| Wild Type | TGGCAGCCTGTCATTGATTACATTGACAGTAAATTTGAAGACTACCTAAATGCAGAATCACGAGTCAACAGACGTCAG  W Q P V I D Y I D S K F E E Y L N A E S R V N R R Q |
| --- | --- |
| Mutant #1 | TGGCAGCCTGTCATTGATTACATT--------------------TGAAGACTACCTAAATGCAGAATCACGAGTCAACAGACGTCAG  W Q P V I D Y I stop |
| Mutant #2 | TGGCAGCCTGTCATTGATT---------------------------- TGAAGACTACCTAAATGCAGAATCACGAGTCAACAGACGTCAG  W Q P V I D L K T T stop |
| Mutant #3 | TGGCAGCCTGTCCTTGATTTGATGGCCAG- AAATTTGAAGATTACCTAAATGCAGAATCACGAGTCAACAGACGTCAG  W Q P V L D L M A R N L K I T stop |

**Supplemental Table 1. Effect of Septin7 genome editing**. Genomic PCR and sequencing were performed to demonstrate the effect of Cas12a mediated genome editing at the Septin7 coding sequence. Three different mutations were detected. The guide RNA sequence is colored in blue. All three mutations lead to premature stop of translation.

**Supplemental Figure S1. Septin inhibitor forchlorfenuron inhibits septin filament assembly specifically.** (A) 100µM of forchlorfenuron (FCF) or the same volume of its solvent ethanol were added to the neural crest explant culture, and the effect on septin filament assembly was analyzed by immunohistochemistry against Septin7. FCF effectively inhibits the formation of long septin filaments in neural crest cells, while EtOH has no effect on septin filament assembly or the spreading of neural crest explants (B). Scale bar = 10µM.

**Supplemental Figure S2.** Septin inhibition reduced focal adhesion size as reflected by GFP-FAK. Live imaging of GFP-FAK expressed in control and FCF treated neural crest cells. Scale bar = 10µm.

**Supplemental Figure S3. Cdc42ep1 misexpression affected the assembly of septin filaments, which secondarily disrupted the formation of actin stress fibers.** A) The organization of septin filaments and actin filaments were defective upon Cdc42ep1 misexpression. B) Higher magnification images of the boxed area in (A). Scale bar = 10µm.

**Supplemental Figure S4:** **The pipeline for quantitative analysis of septin and Cdc42ep1 structures.** A) Segmentation was performed separately for the cells (light blue) and the septin/Cdc42ep1 structures (red). Local background subtraction was performed before thresholding to account for both bright and dim structures. B) Each identified structure was characterized with a set of 70 features including 10 features calculated with MATLAB function (regionprops), the phase invariant form of the first 30 Fourier modes of the border coordinates (shape metrics), and the phase invariant form of the first 30 Fourier modes of the nearest-neighbor distance along the object’s border (relative positioning metrics). C) The first 17 principal components explaining 90% of the variance were used for further analysis. Clustering was performed with DBSCAN algorithm using ε = 2 and the minimal number of points for the core identification = 3. The seven (for septin data) and eight (for Cdc42ep1 data) largest clusters represented 99% of all data points. D) An optional step of cluster refinement was used for five largest clusters in Cdc42ep1 data. Refinement was performed with MATLAB’s Gaussian Mixture Model (GMM) method with the default parameters based on one of the principal component projections (left panel). Here the refinement allowed to separate better elongated and round objects (right panel).

**Supplemental Figure S5:** **The pipeline for quantitative analysis of septin and Cdc42ep1 structures (continued).** A) Image-level quantification was performed based on the area fraction of structures, i.e., the ratio of the total area of structures in one class to the total area of all structures in the image. B) For a more intuitive interpretation of the results, all classes were merged into three larger classes (red, green and blue ellipsoids) based on the biplot showing co-alignment of the unit vector projections of the image-level characteristics along their first two principal components. C) Phenotypic distinction was evaluated based on the area fraction of structures in the three identified classes.

**Supplemental Movie 1. Reassembly of actin stress fibers in control neural crest cells**. Actin filaments in neural crest cells were eliminated by Latrunculin A treatment. Then, Latrunculin A was washed away, and actin reassembly was recorded at 2-min intervals for 1 hour.

**Supplemental Movie 2. Reassembly of actin stress fibers in FCF-treated neural crest cells.** Actin filaments in neural crest cells were eliminated by Latrunculin A treatment. Then, Latrunculin A was replaced by FCF, and actin reassembly was recorded at 2-min intervals for 1 hour.

**Supplemental Movie 3. Trajectories of control explanted neural crest cell migration.** Recorded at 5-min intervals for 4 hours.

**Supplemental Movie 4. Trajectories of Septin7 knockdown neural crest cell migration.** 10ng of Sept7-MO per embryo was injected into neural crest forming blastomeres at cleavage stages. Recorded at 5-min intervals for 4 hours.

**Supplemental Movie 5. Trajectories of Cdc42ep1 knockdown neural crest cell migration.** 10ng of Cdc42ep1-MO per embryo was injected into neural crest forming blastomeres at cleavage stages. Recorded at 5-min intervals for 4 hours.

**Supplemental Movie 6. Trajectories of Cdc42ep1 Septin7 double knockdown neural crest cell migration.** 5ng of Cdc42ep1-MO and 5ng of Septin7-MO per embryo was injected into neural crest forming blastomeres at cleavage stages. Recorded at 5-min intervals for 4 hours.

**Supplemental Movie 7. Dynamics of actin filaments in control neural crest cell migration.** Recorded at 2-min intervals for 1 hour.

**Supplemental Movie 8. Dynamics of actin filaments in neural crest cell receiving 10ng of Septin7-MO.** Recorded at 2-min intervals for 1 hour.

**Supplemental Movie 9. Dynamics of actin filaments in neural crest cell receiving 10ng of Cdc42ep1-MO.** Recorded at 2-min intervals for 1 hour.

**Supplemental Movie 10. Dynamics of actin filaments in neural crest cell receiving 5ng of Septin7-MO and 5ng of Cdc42ep1-MO.** Recorded at 2-min intervals for 1 hour.
