## Supplementary figures and images for "Coordinated Regulation of Cdc42ep1, Actin, and Septin Filaments during Neural Crest Cell Migration"

### Supplemental Figure 1

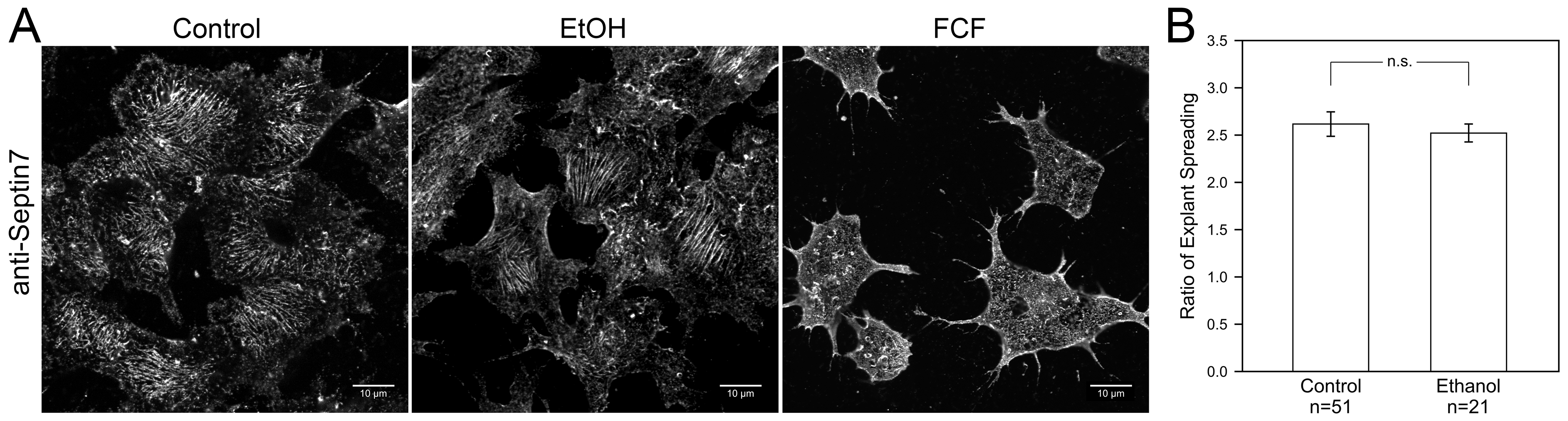

### Supplemental Figure 2

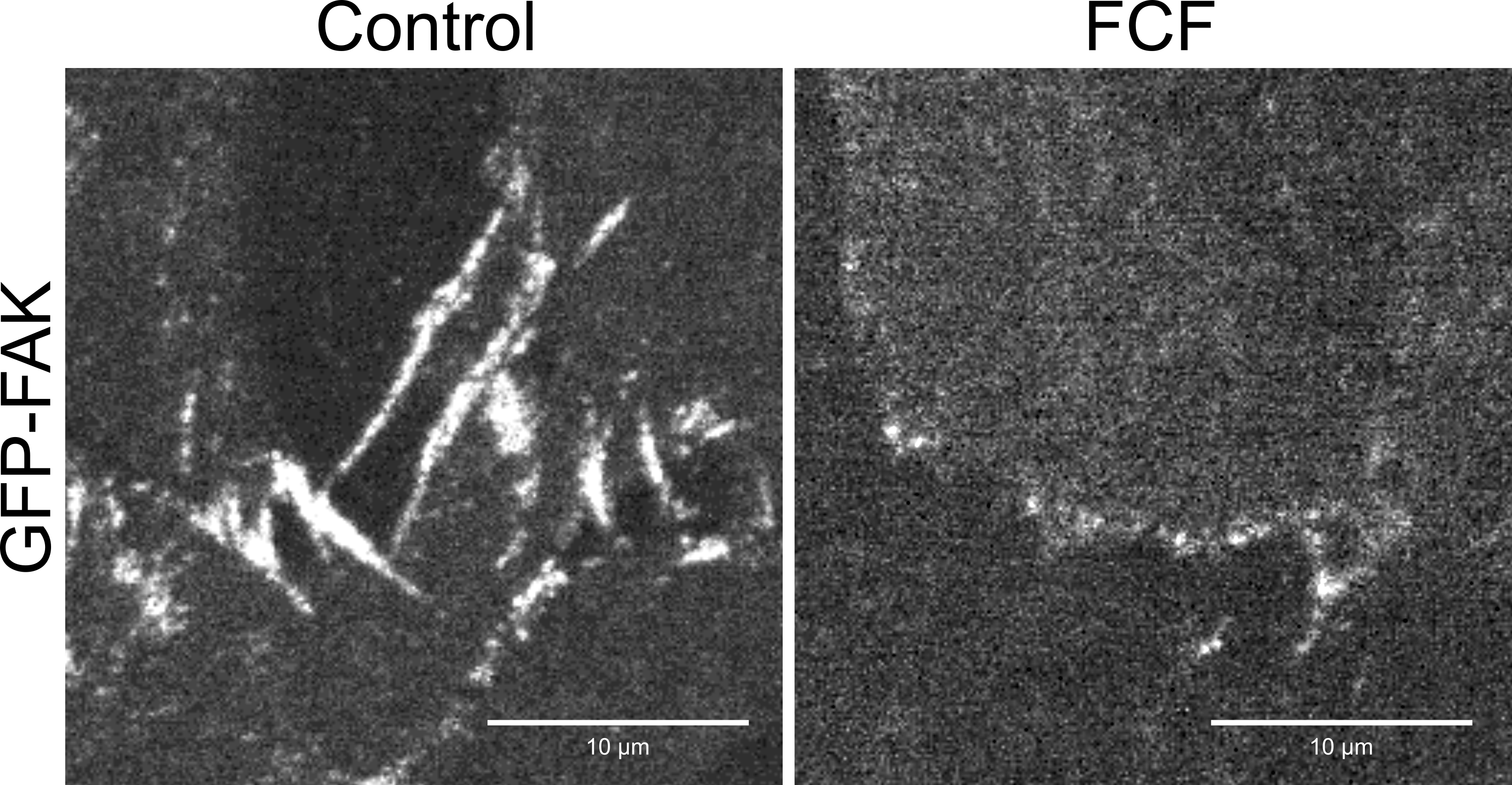

### Supplemental Figure 3

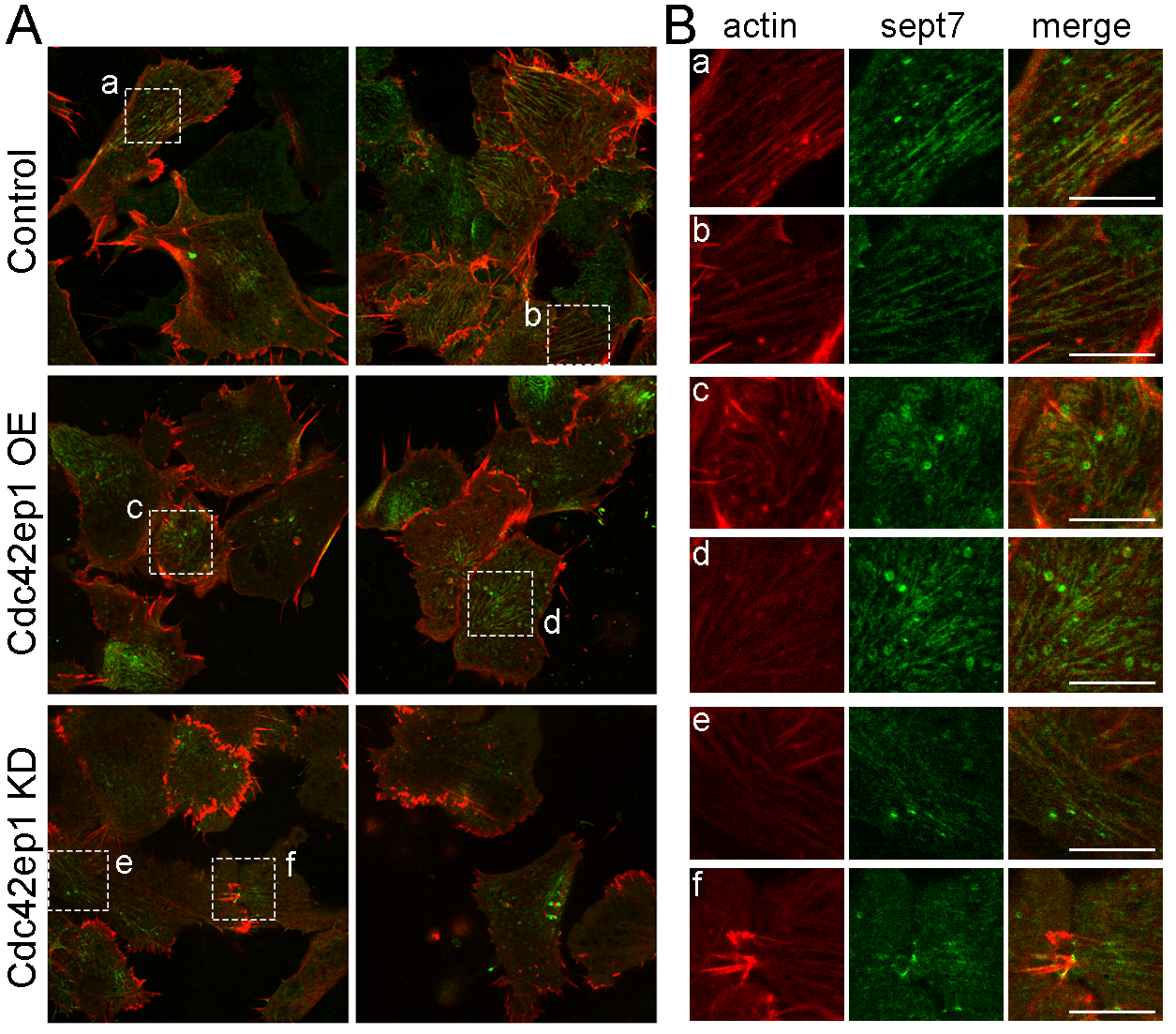

### Supplemental Figure 4

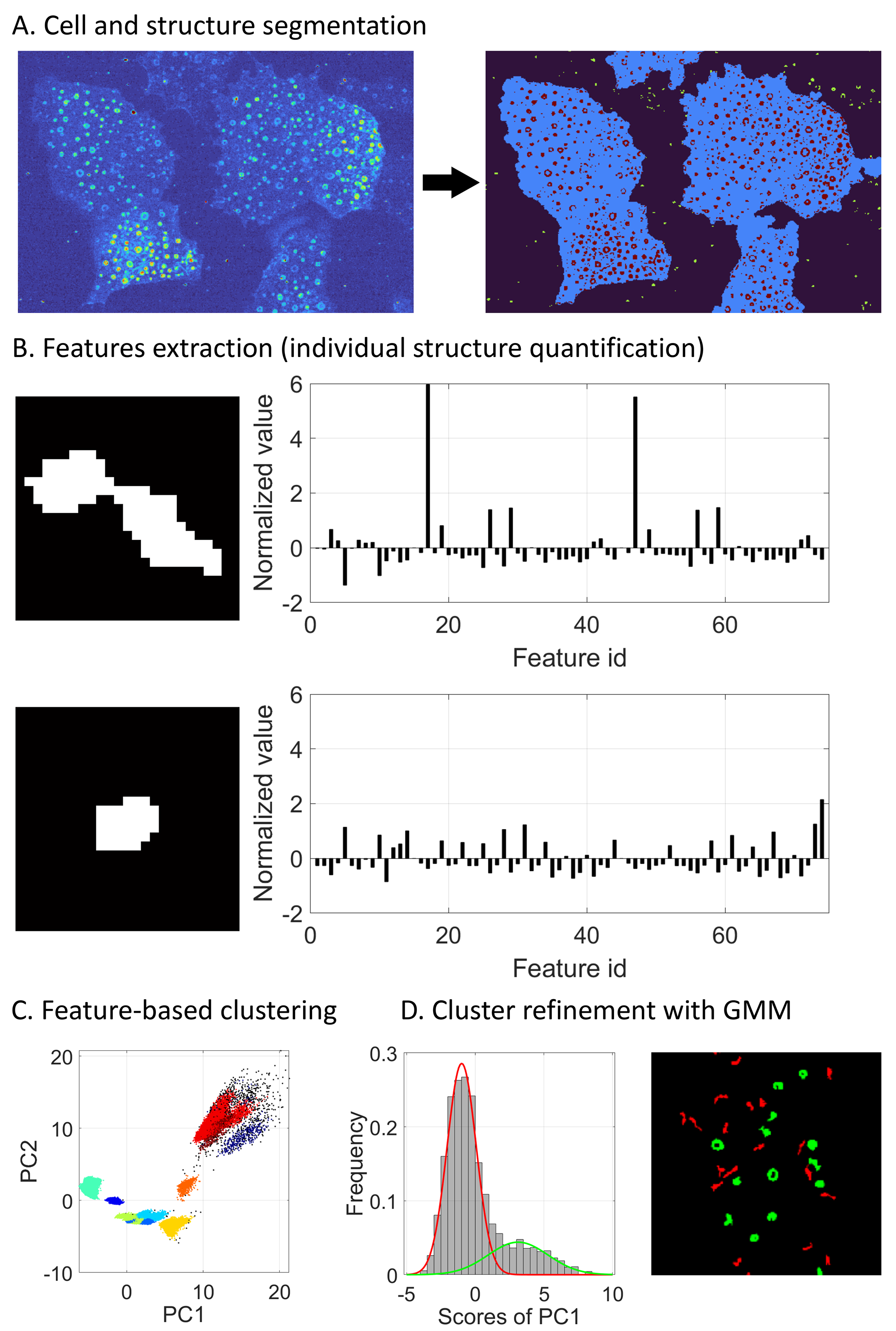

### Supplemental Figure 5

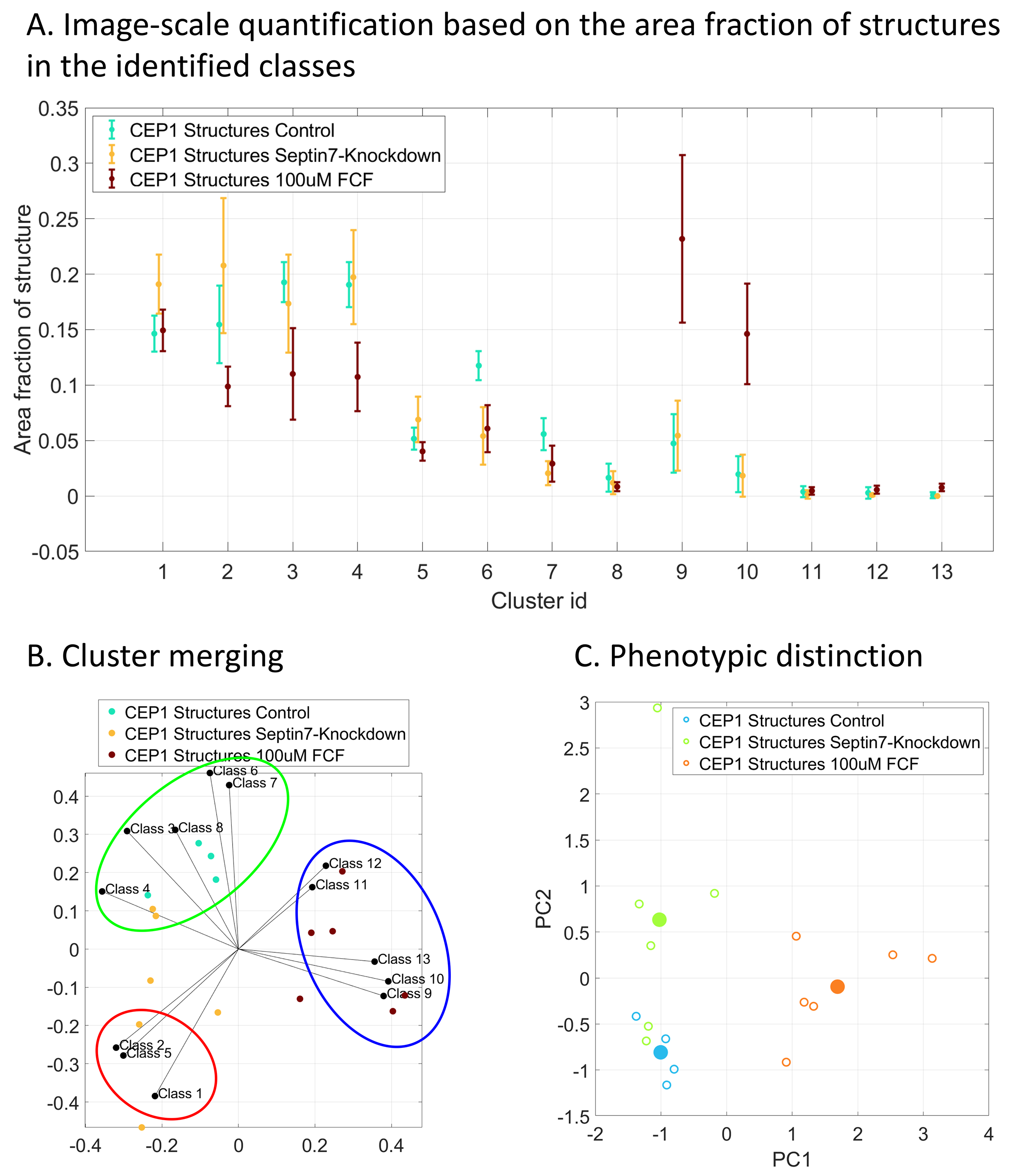
